# Standing on elevated platform changes postural responses during arm movement

**DOI:** 10.1101/2020.08.13.250266

**Authors:** Luis Mochizuki, Juliana Pennone, Aline Bigongiari, Renata Garrido Cosme, Monique Oliveira Baptista Cajueiro, Alberto Carlos Amadio

## Abstract

This study investigated the muscle activity during the preparatory (anticipatory postural adjustment, APA), execution (online postural adjustments, OPA), and compensatory (compensatory postural adjustment, CPA) phases during standing with eyes opened or closed on an elevated platform. Eight healthy young women stood in the upright position, with eyes opened or closed, and did as-fast-as-they-could shoulder flexions on the ground and on 1-m-height-portable-elevated-platform. The surface EMG of trunk (lumbar extensor, and rectus abdominis) and lower limb (rectus femoris, biceps femoris, tibialis anterior and gastrocnemius lateralis) muscles during this task were recorded (1 kHz sampling frequency) and compared during these three phases. Analysis of variance was applied to compare the effects of height (floor and elevated platform), vision (open and closed), and postural adjustment (APA, OPA and CPA) into the activity of each muscle. These muscles were more active during OPA (p<0.0001) and less active during APA. On the elevated platform, these postural muscles presented more activty during APA (p<0.001). During the most stable condition (on the ground with eyes opened), muscle activity during APA and OPA was negatively correlated, and not correlated between OPA and CPA. Our results suggest postural control adapts to sensory, motor, and cognitive conditions. Therefore, the increased demand for postural control, generated due to the height of the support base, provokes the need for greater flexibility of postural synergies and causes a change in muscle activity.

**Summary Statement:** We discuss how postural muscle activity behaves before and after a fast upper arms movement when someone stands on a elevated platform or on the ground.

## Introduction

Anticipation is a cognitive activity based on experience. The predicted outcomes are the key to select the proper action (Schiffer et al., 2015). The anticipatory postural adjustment (APA) is a postural action to ensure the proper mechanical condition for the motor action, for example, to perform daily life activities without falling. Belen’kii et al. (1967) have proposed muscles are active during APA for balance control with the minimum energy expenditure. If the postural muscular activity during APA does not minimize a postural disturbance, compensatory strategies are necessary to ensure system stability (Chabran et al., 1999; Toussaint et al., 1998). Thus, APA (Hay and Redon, 1999) influences the compensatory postural adjustment (CPA), because the central nervous system (CNS) integrates afferent and efferent information to trigger and modulate APA and CPA (Massion, 1998; Maurer and Peterka, 2005; Woollacott and Shumway-Cook, 2002). APA can be modulated by perception (Slijper et al., 2002) and by the motor action itself (Shiratori and Aruin, 2007). Aruin *et al*. (1998) found an APA suppression with high stability demand in an equilibrium recovery task, and this strategy might minimize some instability effects created by the APA itself (Aruin et al., 1998). Nouillout *et al*. (2000) suggest the absence of APA when the configuration of the support base is unstable for equilibrium (Nouillot et al., 2000). These evidences suggest the anticipatory control is conservative.

Predictions are based on perceptions. Non-realistic predictions can emerge from wrong assumptions built from misinterpretations. For example, standing on an elevated platform may induce the fear of falling, although no real physical hazard would be evidente (Adkin et al., 2002). Can such a misinterpretation of falling risk alter the postural actions? The CPA’s interpretation depends on its epoch and it usually does not consider the perturbation end (Bigongiari et al., 2011; Chikh et al., 2018; Kurtzer, 2015), or its nature (collision, De Azevedo et al., 2016; the focal movement, Bigongiari et al., 2011; cyclic movement, (Chikh et al., 2018). These reactive postural actions have a short, medium, and long-latency responses (Kurtzer, 2015). While the long latency component is a transcortical loop contributing to the postural response, the short and medium responses are based on reflex response and voluntary reactions (Santos et al., 2010a). The CPA has different meanings according to the perturbations’ nature. In a collision or any fast perturbation, the perturbation is discrete; however, the perturbation can be a transient. Focal movement can be a postural perturbation transient, which is a sequence of two discrete perturbations (when the focal movement switches on and off). The standard CPA emerges when focal movement switches on. When the focal movement switches off, another reactive response is induced by the follow-through phase. The follow-through is usually known as the final phase in baseball pitching (Chu et al., 2016), golf swing (Steele et al., 2018), and basketball shooting (Okazaki et al., 2015). To differentiate these two reactive postural actions, we suggest calling the online postural adjustments (OPA) the postural action during the focal movement, while we will call CPA as the postural response during the follow-through. In APA studies (Belenkii, 1967; Yiou, 2012), although the arm raising is a discrete movement, no study has concerned about the follow-through phase. The perception of these two discrete perturbations might trigger two different postural reactions.

Postural responses are scaled by perception. Adkin et al. (2000) showed postural threat modulates the postural control. Psychological factors impact perception and balance. For example, anxiety generates balance disturbances (Sturnieks et al., 2016). Persons can feel less postural confidence standing on high places, and under this condition, closing the eyes might impair the ability to maintain postural stability (Aruin et al., 2001; Maki et al., 1990; Palm et al., 2009; Paulus et al., 1984; Vuillerme et al., 2001). Such confidence loss might lead to an avoidance behavior, such as the fear of falling. Fear of falling manifests by refusing to stand still in high places with a narrow support base, or by the elderly refusing to move after an accidental fall (Adkin et al., 2000; Maki et al., 1990; Yiou, 2012). Fear of falling affects balance control (Adkin et al., 2002; Maki et al., 1990; Sartor-Glittenberg et al., 2018; Sturnieks et al., 2016), and APA is reduced in amplitude e elongated in time (Adkin et al., 2002; Maki et al., 1990; Tisserand et al., 2016; Yiou, 2012). Although APA does not fully attenuate the postural perturbation, few studies attempted to describe the postural adjustments occurring during and after the focal movement (Bouisset et al., 2000; Le Bozec et al., 2008; Santos et al., 2010b) under circumstances that the fear of falling can emerge. We suggest this non-realistic postural threat may lead to a perception misinterpretation and change the postural actions.

To describe all attempts to stabilize the body due to the focal movement’s postural perturbations, this study investigated the postural adjustments’ behavior along the whole action, in the preparatory (APA), execution (online postural adjustments, OPA), and compensatory (CPA) phases during standing with eyes opened or closed. This study aims to verify the effects of reduced sensorial information and different heights on postural muscles activity during these three phases. Our main hypothesis is standing on high ground will induce changes in muscle activity along the action. To reach this aim, young women did as-fast-as-possible shoulder flexion while standing on the ground and on 1-m-high-portable-elevated-platform. Compared to other studies about APA and CPA, this is a different temporal division for the postural responses’ organization, because postural activity during the follow-through phase and APA was rarely studied. Considering the focal movement triggers two perturbations (when it begins and when it ends), the postural muscle activity during OPA might affect what comes before (APA) and after (CPA). Our second hypothesis the postural muscles activity during the focal movement will be higher compared with APA or CPA, and it will be enhanced by the elevated ground. Thus, we expected postural activity during focal movement would modulate the postural muscle activity during APA and CPA. Therefore, the analysis of OPA would give us more information about how postural control deals with such an avoidance behavior. We have also calculated the postural muscle synergies during these phases and discussed the association between such synergies and the focal movement.

## Method

### Participants

Students at the University campus were invited to attend this study. Participants were eight healthy young women (22±3 years old, 1,59±0,05 m tall and 58.7±4.2 kg mass). The inclusion criteria were not to have any neurological or musculoskeletal injury or disorder, and not to have any balance disorder. Participants would be excluded if they were not able to stand and feel safe on the platform (1 m high, 0.5 m side). All participants signed an informed consent term in accordance with the Ethical Committee of the University of São Judas Tadeu.

### Instruments

The muscular electrical activity (EMG) and the shoulder angle were measured. Data sampling frequency was 1 kHz. A two-dimensional flexible electrogoniometer (NorAngle II, Noraxon, Inc USA) was used to measure the shoulder motion. The EMG was recorded with 8-channel EMG system (Myosystem 1400, Noraxon, Inc, USA; 20 to 500 Hz band-pass filter; input impedance >10 MΩ; Common Rejection Mode >85 dB; Noise Ratio <1μV RMS; Total amplification 1000 times) using active (differential and pre-amplified) surface electrodes. These are the selected muscles: anterior deltoid (AD), lumbar extensor (LE), rectus abdominis (RA), rectus femoris (RF), biceps femoris (BF), tibialis anterior (TA) and gastrocnemius lateralis (GL). The electrodes’ placement followed the SENIAN procedures (Hermens et al., 2000) with 1 cm between electrode centers. These two measurement systems are connected to a data acquisition system controlled by the Myoresearch software installed in PC Pentium 4 2.66 MHz computer.

### Motor Task

In a standing position (feet parallel, ankles and haluxes touching), the task was to raise both arms as fast as possible with extended elbows by flexing the shoulders, holding one 2-kg-barbell in each hand, and suddenly end this movement when both arms were parallel to the ground. The initial position was quiet standing with the arms losing by the trunk’s side. The final position was standing with arms parallel to the ground. This motor task was performed with eyes opened and closed and standing on the ground and standing on 1-m-height platform (0.5 m side). When eyes were closed, they remained like this throughout all the execution, even in the resting position. Participants should keep staring or facing a 0.5 m radius circular target on the wall in front of them 2.5 m ahead, move their arms as fast as possible, and hold the final position for approximately 2 s. After the task was over, they returned to the rest position and hold it for 3 s until the next repetition. The task was self-initiated. Each participant did 4 sets of 10 repetitions. To avoid a reaction-time-like task, we told them to start the task whenever they wanted, but they should hold on for at least 3 seconds their arms in the rest position and in the final position; and always should return to initial position before to start another repetition. Between each series there was a rest interval of 3 minutes to avoid the effects of fatigue. The order of conditions was randomly assigned.

### Variables and data analysis

The raw EMG signal had the mean removed, low-pass filtered with 4th order Butterworth filter with 200 Hz cut-off frequency, and full-wave rectified. The shoulder joint angle signal was low pass filtered with 4th order Butterworth filter with 20 Hz cut- off frequency. For each trial, data (processed EMG and shoulder angle) were cropped into three epochs: APA, OPA, CPA. The initial t_0_ and final t_f_ of shoulder movement, indicated by the onset of the angular acceleration of the shoulder, obtained by double derivation of the angular position measured by the electrogoniometer, were used to define the epochs. The limits of each epoch were: APA (from 200 ms before t_0_ to 50 ms after t_0_); OPA (from 50 ms after t_0_ to t_f_); and CPA (from t_f_ to 250 after t_f_). The APA limits t_1_ and t_2_ vary in the literature (t_1_ from −200 ms to −100 ms, and t_2 from_ 0 ms to 50 ms (Aruin and Latash, 1995; Chen and Tsai, 2016; Krishnan et al., 2012). Then, for APA and the other epochs, we chose the limits t_1_ and t_2_ to have the largest epoch to preserve data information.

The set of processed EMG signals in each postural adjustment was transformed into the EMG principal components (EMG-PC) using the principal component analysis (PCA). PCA is a linear transformation to calculate the eigenvectors and eigenvalues of data matrix (Molenaar, Wang, 2013). We applied the PCA to calculate, from the same set of processed EMG, two, three or four EMG-PC. We did such variation to evaluate the effect of dimension reduction within the synergy. For independent synergies, we took the EMG-PC and applied the independent component analysis (ICA) (Makeig, Westerfield, Enghoff, Jung, Townsend, Courchesne, 2002). The independent components (IC) are statistically independent; thus, we called as muscle synergy the EMG-IC (Mochizuki et al., 2006). All data processing and time series analysis were done with Matlab (version 2009b; Mathworks, USA) scripts.

From the processed EMG of each muscle, the Root Mean Square (RMS) value was calculated for the APA, OPA and APC epochs. The following parameters were calculated in APA, OPA and CPA. To separate these parameters, we use Winter’s proposal (Winter, 1995) on the levels of organization of the neuro-musculoskeletal-joint integration for the movements’ execution.

The parameters of the basic mechanisms of neuromuscular regulation: 1) EMG intensity:

1. RMS value normalized by 95% of the maximum RMS value measured during repetition.
2. Muscle latency: time between the beginning of muscular activation and the beginning of focal movement. The latency of each muscle before the onset of focal movement was calculated. The onset of muscle activity was the first EMG signal, before the FM, three times greater than the baseline signal plus 3 standard deviations of the EMG measured in the pre-activity phase.
3. R index (equation 1): module of the difference between the EMG intensity of the agonist and antagonist muscle of the referred joint. This parameter represents the reciprocal inhibition parameter.

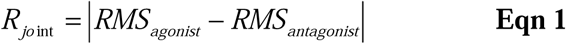
4. C index (equation 2): module of the sum of the EMG intensity of the agonist plus antagonist muscle of the referred joint. This parameter represents the coactivation parameter.

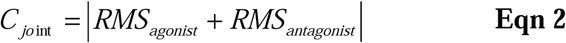 Indicator of the postural synergistic action:
5. Synergy: s*e*t of *n* independent components from the processed EMG (EMG- IC*n*).
6. Synergy variability – relative participation of each PC in the total variance of the EMG-PC*n*.

### Statistical Analysis

For the statistical analysis, analysis of variance, ANOVA were applied assuming 5% as the significant level. We did different comparisons: a) the RMS of each muscle across muscles, conditions (height of support base and vision) and types of postural adjustments; b) the muscle latency across muscles and conditions; c) R and C indexes across joints, conditions and types of postural adjustments; d) the accounted variability of the first three PC across conditions and postural adjustments; and e) the minimum number of principal components to reach at least 75% of the total variance across the conditions and postural adjustments. Tukey HSD post hoc test was used to identify the differences within the factors. For the multiple linear regression, we compared the RMS of residuals across the dimensions and postural adjustments. Statistical analysis was performed using Statistica (version 5.1, 1996, USA).

## Results

Muscle activity (Table 1) was not affected by the support basis height. The activity of all muscles was affected by Postural Adjustment (F_2,1035_>63, p<0.0001). The *post hoc* Tukey HSD test indicated the highest activity of all muscles occurred durig OPA, and the lowest occurred during APA (except AD) or CPA (p<0.0001). The interaction between Postural Adjustment and support base height affected the activity of DA, RA, RF and GL muscles (F_2,1035_>4.7, p<0.008). The *post hoc* test showed AD muscle had the greatest activity in APA and on the ground (except for OPA); RA were the most active on the ground in all postural adjustments; RF presented the highest activity at the top; and GL presented the highest activity at the top in all Postural Adjustments (p<0.001). BF muscle activity was affected by visual information (F_1,1035_=3.9, p<0.05). The *post hoc* Tukey HSD test showed the highest activity occurred when the task was performed with the eyes closed (p=0.02).

**Table 1.** Mean ± Standart Deviation of muscle activity (anterior deltoid AD, lumbar extensor LE, rectus abdominis RA, rectus femoris RF, biceps femoris BF, tibialis anterior TA, gastrocnemius lateralis GL) amplitude (%MAXIMUM) according to the height of support base (floor and elevated platform), vision (open and closed) and Postural Adjustments conditions (anticipatory postural adjustement, APA; online postural adjustment, OPA; compensatory postural adjustment, CPA). OPA* indicates the muscle activity was the highest during OPA compared with APA and CPA (p<0.0001).

| Height | Vision | Postural Adjustment | AD | LE | RA | RF | BF | TA | GL |
| --- | --- | --- | --- | --- | --- | --- | --- | --- | --- |
| Floor | Open eyes | APA (n=80) | 56.0±21.8 | 35.0±10.8 | 15.6±6.6 | 47.7±11.4 | 52.5±16.7 | 1.7±1.2 | 35.5±29.7 |
| | | OPA* (n=80) | 66.4±18.1 | 73.6±15.6 | 71.4±8.3 | 65.9±8.1 | 66.8±11.6 | 78.3±9.8 | 73.4±12.6 |
| | | CPA (n=80) | 48.3±15.4 | 57.2±15.5 | 62.1±10.1 | 55.5±10.6 | 56.7±16.7 | 47.0±10.9 | 46.4±10.7 |
| | Closed eyes | APA (n=80) | 63.6±31.3 | 35.1±11.0 | 15.2±8.0 | 54.4±45.8 | 54.6±14.9 | 1.7±1.1 | 30.9±27.4 |
| | | OPA* (n=80) | 66.7±19.3 | 73.6±15.6 | 70.2±7.4 | 67.3±10.3 | 65.0±11.7 | 76.4±11.0 | 70.8±11.0 |
| | | CPA* (n=80) | 46.3±16.7 | 59.2±16.6 | 62.3±8.8 | 54.8±11.2 | 53.5±14.3 | 46.5±12.6 | 48.2±12.0 |
| Elevated platform | Open eyes | APA (n=80) | 51.2±18.0 | 35.2±9.3 | 19.8±9.5 | 49.9±11.5 | 54.1±15.4 | 1.8±1.2 | 29.8±25.2 |
| | | OPA* (n=80) | 65.3±15.7 | 66.5±8.8 | 70.1±9.4 | 63.5±8.5 | 66.6±10.8 | 76.4±10.8 | 84.9±21.1 |
| | | CPA (n=80) | 52.5±13.5 | 57.8±12.3 | 59.5±9.3 | 55.6±11.1 | 56.3±14.2 | 50.3±12.4 | 47.4±10.1 |
| | Closed eyes | APA (n=80) | 52.4±28.2 | 35.0±8.7 | 17.8±7.6 | 51.3±12.3 | 54.2±16.6 | 2.4±2.3 | 29.4±25.4 |
| | | OPA* (n=80) | 67.2±14.6 | 71.2±12.2 | 72.3±8.5 | 67.2±15.6 | 66.9±12.5 | 77.9±10.5 | 73.2±13.4 |
| | | CPA (n=80) | 52.1±17.0 | 60.1±15.0 | 61.7±8.2 | 57.1±11.1 | 55.2±13.4 | 47.9±12.0 | 51.1±10.9 |

Correlation analysis was performed to associate the muscle activities between APA and OPA, between OPA and CPA, and between APA and CPA. For APA and OPA, the muscle activities were correlated for all conditions [ground and eyes opened (R^2^=0.64 p=0.02), ground and eyes closed (R^2^=0.63 p=0.02), elevated and eyes closed (R^2^=0.88 p<0.001)], but when participants were with eyes opened on the elevated platform (R^2^=0.18 p=0.19). For APA and CPA, muscle activities were not correlated for all conditions [ground and eyes opened (R^2^=0.11 p=0.54), ground and eyes closed (R^2^=0.15 p=0.65), elevated and eyes opened (R^2^=0.05 p=0.43), elevated and eyes closed (R^2^=0.02 p=0.40)]. For OPA and CPA, the muscle activities were not correlated for all conditions [ground and eyes opened (R^2^=0.21 p=0.17), ground and eyes closed (R^2^=0.10 p=0.54), elevated and eyes closed (R^2^=0.22 p=0.16)], but when participants were with eyes opened on the elevated platform (R^2^=0.65 p=0.02).

The R index is presented in Table 2. The R index was affected by the support base height (F_1,1025_=22.6, p<0.0001), postural adjustment (F_2,1035_=92.7, p<0.0001) and joints (F_2,2070_=92.7, p<0.0001). The *post hoc* Tukey HSD test showed R index was the highest on the ground, in APA and at the ankle (p<0.0001), and the lowest in OPA and at the knee (p<0.0001).

**Table 2.** Mean ± Standart Deviation of R index (%MAXIMUM) according to the height of support base (floor and elevated platform), vision (open and closed) and Postural Adjustments conditions (anticipatory postural adjustement, APA; online postural adjustment, OPA; compensatory postural adjustment, CPA). Floor* indicates R index was higher when participants did the task on the floor than on the elevated plataform (p<0.0001). APA* indicates R index was higher during APA than during OPA or CPA. R-ankle* indicates R index was higher for the ankle joint than for the hip and knee joints (p<0.0001).

| Height | Vision | Postural Adjustment | R-HIP | R-KNEE | R-ANKLE* |
| --- | --- | --- | --- | --- | --- |
| **Floor\*** | **Open eyes** | **APA\* (n=80)** | 19.4±10.1 | 17.8±14.2 | 33.9±29.5 |
| | | **OPA (n=80)** | 14.6±12.8 | 12.7±8.3 | 13.0±12.3 |
| | | **CPA (n=80)** | 16.7±11.7 | 15.9±11.6 | 13.7±10.2 |
| | **Closed eyes** | **APA\* (n=80)** | 20.3±10.0 | 16.4±42.4 | 29.2±27.2 |
| | | **OPA (n=80)** | 13.5±11.9 | 11.2±7.5 | 14.2±10.6 |
| | | **CPA (n=80)** | 14.3±11.9 | 13.0±9.2 | 17.1±11.0 |
| **Elevated Platform** | **Open eyes** | **APA\* (n=80)** | 16.4±11.3 | 10.9±8.1 | 28.0±24.9 |
| | | **OPA (n=80)** | 8.9±7.7 | 10.9±9.1 | 13.0±15.8 |
| | | **CPA (n=80)** | 11.5±7.9 | 11.1±9.2 | 12.8±9.2 |
| | **Closed eyes** | **APA\* (n=80)** | 17.6±9.9 | 14.1±10.2 | 27.2±25.0 |
| | | **OPA (n=80)** | 11.9±10.7 | 11.7±15.9 | 12.5±12.1 |
| | | **CPA (n=80)** | 12.8±10.5 | 12.5±9.2 | 16.2±10.9 |

The C index is presented in Table 3. The C index was affected by Postural Adjustment (F_2,1035_=3313, p<0.0001), and joints (F_2,2070_=339, p<0.0001). The Tukey HSD *post hoc* test showed C index was the highest in OPA (p<0.0001), and at the knee (p<0.0001), and the lowest in APA and at the ankle (p<0.0001).

**Table 3.** Mean ± Standart Deviation of C index (%MAXIMUM) according to the height of support base (floor and elevated platform), vision (open and closed) and Postural Adjustments conditions (anticipatory postural adjustement, APA; online postural adjustment, OPA; compensatory postural adjustment, CPA). OPA* indicates C index was higher during OPA than during APA or CPA (p<0.0001). R-knee* indicates R index was higher for the knee joint than for the hip and ankle joints (p<0.0001). APA” indicates R index was lower during APA than during OPA or CPA (p<0.0001). R-ankle” indicates R index was lower for the ankle joint than for the hip and knee joints (p<0.0001).

| Height | Vision | Postural Adjustment | C-HIP | C-KNEE* | C-ANKLE'' |
| --- | --- | --- | --- | --- | --- |
| Floor | Open eyes | APA'' (n=80) | 50.7±14.7 | 100.2±17.8 | 37.2±29.8 |
| | | OPA* (n=80) | 145.0±15.8 | 132.7±13.1 | 151.7±14.5 |
| | | CPA (n=80) | 119.3±17.1 | 112.1±19.9 | 93.5±13.2 |
| | Closed eyes | APA (n=80) | 50.3±15.8 | 109.1±50.6 | 32.6±27.7 |
| | | OPA* (n=80) | 143.8±16.9 | 132.4±17.5 | 147.2±14.1 |
| | | CPA (n=80) | 121.5±19.1 | 108.2±20.2 | 94.7±14.0 |
| Elevated Platform | Open eyes | APA (n=80) | 55.0±13.9 | 104.0±23.8 | 31.6±25.6 |
| | | OPA* (n=80) | 136.6±14.3 | 130.1±13.6 | 161.4±27.8 |
| | | CPA (n=80) | 117.3±16.8 | 111.9±21.0 | 97.7±16.4 |
| | Closed eyes | APA (n=80) | 52.7±12.5 | 105.5±23.6 | 31.8±25.7 |
| | | OPA* (n=80) | 143.5±13.6 | 134.1±20.3 | 151.1±17.3 |
| | | CPA (n=80) | 121.8±17.7 | 112.4±19.0 | 99.0±12.4 |

### Postural synergies

The first three principal components variances are shown in Table 4. These three principal components were affected by the Postural Adjustment (F_2,1035_>473, p<0.001), and two of them (PC1 and PC3) were affected by the support base height (F_1,1035_>4.4, p<0.04). The *post hoc* Tukey HSD test showed the PC1 explained variance was the highest in APA and at the ground and the lowest in OPA (p<0.0001); the PC2 explained variance was the highest in OPA and the lowest in APA (p<0.0001); and PC3 explained variance was the highest in OPA and at the top and the lowest in APA (p<0.0001).

**Tabela 4.** Mean ± Standart Deviation of principal component accounted variability (Principal component 1, PC1; Principal component 2, PC2, Principal component 3, PC3) according to the height of support base (floor and elevated platform), vision (open and closed) and Postural Adjustments conditions (anticipatory postural adjustement, APA; online postural adjustment, OPA; compensatory postural adjustment, CPA). * indicates the highest value for the comparison between Floor and Elevated platform, and “indicates the lowest value for the comparison between Floor and elevated platform.

| Height | Vision | Postural Adjustment | PC1 (%) | PC2 (%) | PC3 (%) |
| --- | --- | --- | --- | --- | --- |
| Floor | Open eyes | APA (n=80) | 93.4±3.7* | 3.6±2.6” | 1.6±1.0 |
| | | OPA (n=80) | 67.8±7.4” | 12.1±4.3* | 6.8±2.0 |
| | | CPA (n=80) | 78.1±6.5 | 10.2±4.3 | 5.3±1.9 |
| | Closed eyes | APA (n=80) | 93.5±3.6* | 3.7±2.8” | 1.5±0.9 |
| | | OPA (n=80) | 68.5±7.1” | 12.2±4.2* | 6.7±1.8 |
| | | CPA (n=80) | 79.4±5.6 | 9.6±3.3 | 5.0±1.6 |
| Elevated Platform | Open eyes | APA (n=80) | 93.5±3.4 | 3.6±2.5 | 1.5±0.7” |
| | | OPA (n=80) | 66.9±10.3 | 12.8±5.6 | 7.2±2.6* |
| | | CPA (n=80) | 77.2±6.6 | 10.6±4.1 | 5.4±2.2 |
| | Closed eyes | APA (n=80) | 92.9±3.8 | 3.8±2.7 | 1.7±0.8” |
| | | OPA (n=80) | 67.7±8.0 | 12.5±5.1 | 6.7±1.7* |
| | | CPA (n=80) | 76.1±7.1 | 10.7±4.0 | 5.8±2.0 |

## Discussion

In this study, young women did as-fast-as-possible shoulder flexion while standing on the ground and on 1-m-high-portable-elevated-platform. We recorded the EMG of trunk and lower limb muscles during the preparatory, execution and compensatory postural phases. Although average muscle activity was similar on the ground and on the elevated platform, how postural muscles were active across the motor action was different. All postural muscles presented the highest activity during OPA, and the lowest in APA. Standing on an elevated platform changes how trunk and lower limb muscles are activated during an upper limb task: RA reduced its activation, while RF and GL has increased their activation. During the most stable condition (on the ground with eyes opened), muscle activity was negatively correlated between APA and OPA, and not correlated between OPA and CPA. These results suggest how flexible the postural activity is to adapt to the sensory constraints and a false threat. Indeed, two facts suggest standing on an elevated platform induces a conservative attitude. Although no mechanical differences exist, standing on an elevated platform induced an increased muscle activity during OPA. Besides, muscle activity during APA reduced when the participants stood on an elevated platform. This conservative attitude suggest that such condition is really a postural threat.

The timeline of muscle activation reveals the plot of postural activities during the motor action. Our results showed that muscle activity was different throughout the reactive postural control phases. Moreover, these results support the idea of dividing the postural muscle activity in three phases. Usually, postural studies have only one compensatory postural phase (Bigongiari et al., 2011; Chen and Tsai, 2016; Chen et al., 2015; De Azevedo et al., 2016; Knox et al., 2016; Labanca et al., 2015; Schlenstedt et al., 2017). For these studies, CPA starts with the execution of the FM. Operationally, we have defined OPA as the activity of postural muscles that occurs during the execution of FM; while, CPA is the activity of the postural muscles that begins with the end of FM up to 250 ms later, the follow-through phase. During OPA, fast corrections, based on reafference control, support the focal movement performance; while feedback control during CPA to concern with any postural problems (balance, body orientation or instability) induced by follow-through. Other studies have suggested different phases for the anticipatory postural adjustments (Krishnan et al., 2012) and for the reactive control. Latash and his colleagues have proposed the early anticipatory postural adjustment and the early anticipatory synergy adjustment (Krishnan et al., 2012) as two important control mechanisms in postural control.

Less confidence due the postural threat changes the relation between APA and OPA. Muscle activity during the focal movement and during the anticipatory postural activity were correlated, but this association was lost with opened eyes. The postural threat induced by the elevated platform is fed by visual information. On the other hand, standing on the elevated platform with opened eyes induced muscle activity to be associated during OPA and CPA. This postural threat induces different associations between muscle activations. On the ground, the focal movement triggers the preparation phase (during APA), but does not tune the postural activity during the compensatory phase. However, when participants were on the elevated platform, a confidence lack has just emerged with closed eyes, and such relations have changed. Although, the focal movement has not triggered muscle activation during the preparatory phase with opened eyes, but during the compensatory phase. Perception might has affected these relations because the task mechanical status remained the same. This result supports the hypothesis that the postural threat induced by the elevated platform would induce changes in muscle activity across the whole motor action”.

Elevated platform inducing a postural threat has changed the organization of postural responses and motor action organization. Comparing the high and low support conditions, RA activation decreased, while RF and GL have increased their activation on the elevated platform. These two biarticular muscles suggest a higher postural demand during the task. Adkin et al. (2002) and Adkin et al. (2003) showed APA activity was inversely associated with the fear of falling (Adkin et al., 2002; Adkin et al., 2003). Both studies have shown their participants did the shoulder flexion on elevated support base or closer to the support base’s edge, less anticipatory activity was observed. In the same way, Gendre et al. (2016) showed the effects of fear of falling depend on initial environmental conditions added on the APA direction relative to the postural threat (Gendre et al., 2016).

Agonist and antagonist muscle activations have different patterns across joints and postural adjustments. Coactivation and reciprocal inhibition indexes have changed across joints and postural adjustments. Modulation of postural control, acting through muscle pairs, can be explained by the changes of these two indexes (Bigongiari et al., 2011; Slijper et al., 2002). These indexes are different between joints and Postural Adjustment. Reciprocal inhibition reduced in the elevated platform, while coactivation remained the same. This is an example of mechanical and perceptual factors affecting balance control. As the mechanical balance conditions were not modified by the elevated platform, the greater coactivation during OPA suggests another effect of fear of falling (Carpenter et al., 2001). Our results showed greater muscle activity in some postural and focal muscles with the high support base, which suggests an increase in voluntary activity to perform the same motor task.

The greatest reciprocal inhibition occurred in the APA and ankle, when the number of dimensions was smaller. The greatest number of dimensions occurred during the OPA, with the highest coactivation, mainly in the knee, and when the task was performed with the elevated support base. These results suggest that the organization of postural adjustments is flexible. Matrix factorization methods are applied to find muscle synergies (Lambert-Shirzad and Van der Loos, 2017) or muscle modes (Krishnan et al., 2012). Muscle synergies and muscle modes indicate how muscles can be clustered in relation to the task. In both cases, the number of muscle synergies or modes indicates the complexity of the system. Thus, our results indicate that antagonistic muscles activate less and facilitate movement in low complexity tasks or activate more and prevents movement in more complex tasks, such as in elevated support base.

The muscle latency was not altered while standing on 1-m-high-portable- elevated-platform. This postural threat is not a mechanical threat since the higher support base was completely fixed and stable. Comparing the principal components during task performed on the ground and on the elevated platform, PC1 have decreased its accounted variability and PC3 increased. The greater PC1 participation when the task was on the ground reinforces its conservative nature, while the increased PC3 participation reinforces its restorative role in postural stability. On the ground, postural control acted with a reduced number of dimensions. While the execution of the focal movement requires the participation of more main components, suggesting the increase of the complexity of its control.

This study presents some limitations that should be noted. First, the small sample size does not support a large statistical power. Although there is no evidence of difference in the neuromuscular system between sexes, our sample may be less representative of the population because it consists only by women. Last, but not least, as there is no mechanical difference between performing the movement on ground or on elevated platform, changes in muscle activity are most likely due to fear of falling, but fear was not measured by any kind of scale of perception.

## Conclusion

The analysis of postural control divided into three phases seems to be interesting to elucidate the strategies used by the CNS to plan and execute a movement. The results showed how postural control, by means of individual muscle activation, agonist and antagonist muscle pairs, muscle synergies, acts along the experimental conditions. These results suggest postural control adapts to sensory, motor, and cognitive conditions. Therefore, the increased demand for postural control, generated due to the height of the support base, provokes the need for greater flexibility of postural synergies and causes a change in muscle activity.

### Practical applications

The fear of falling can directly influence the balance even without any mechanical threat. Therefore, interventions that encourage stabilization may have their complexity increased only by changing the support base with the aim, for example, of improving gait and decreasing the risk of falls in the elderly.

**Figure 1.**
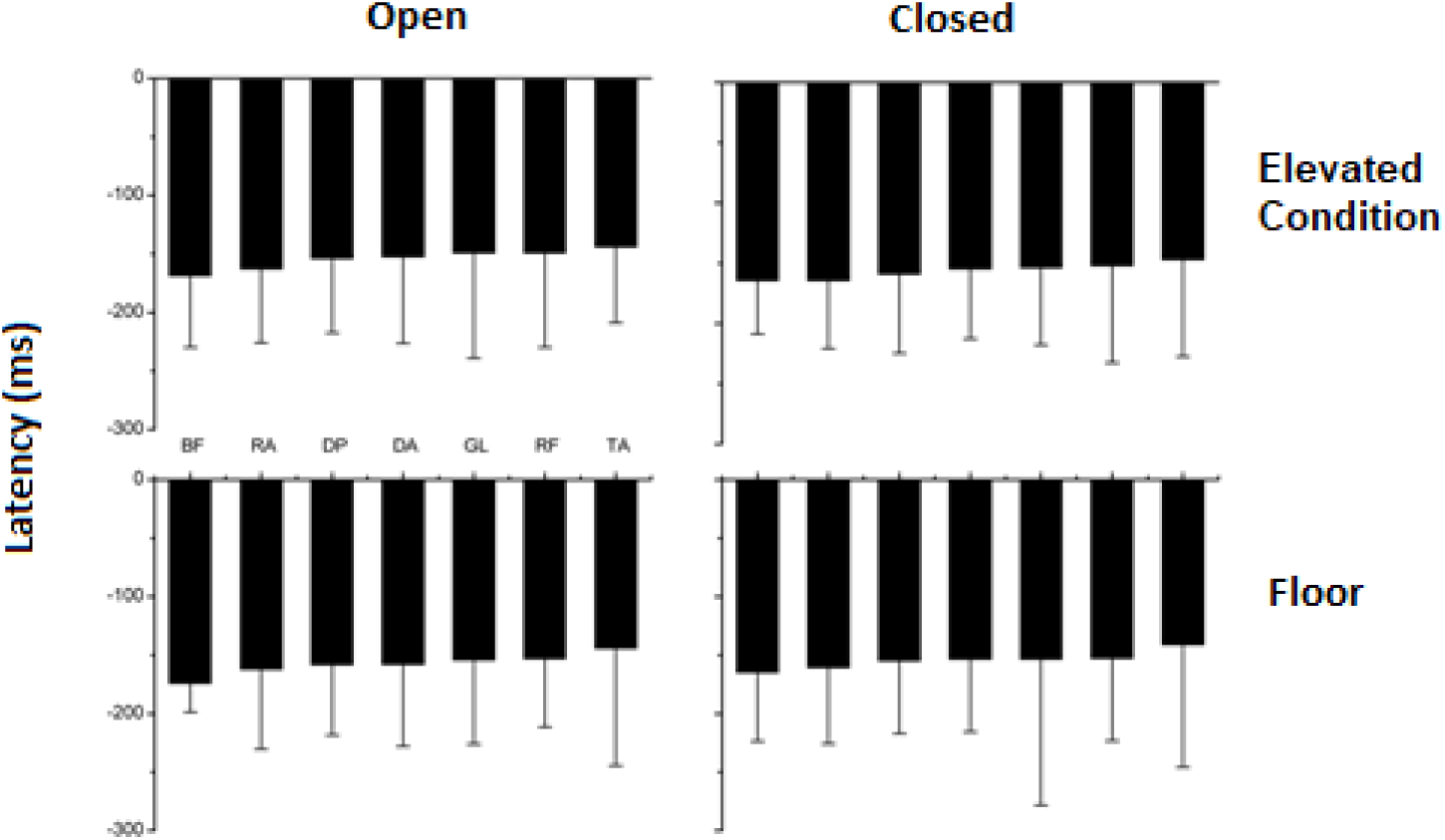
Mean ± Standart Deviation (n=80) of latency (anterior deltoid AD, lumbar extensor LE, rectus abdominis RA, rectus femoris RF, biceps femoris BF, tibialis anterior TA, gastrocnemius lateralis GL) according to the height of support base (floor and elevated platform) and vision (open and closed) conditions.

## Declarations

### Funding (information that explains whether and by whom the research was supported)

This research was supported by National Council for Scientific and Technological Development, Brazil.

### Conflicts of interest/Competing interests (include appropriate disclosures)

Luis Mochizuki, Juliana Pennone, Aline Bigongiari, Renata Garrido Cosme, Monique Oliveira Baptista Cajueiro and Alberto Carlos Amadio declare that they have no conflict of interest.

### Ethics approval (include appropriate approvals or waivers)

School of Arts, Sciences, and Humanities Ethics Committee, project number 59263516.6.0000.5390

### Consent to participate (include appropriate statements)

Participants were informed about this study and they agreed to attend the experimental session. Participants read and signed the informed consent statement.

### Availability of data and material (data transparency)

Not applicable.

### Code availability (software application or custom code)

No applicable.

